# Strain- and age-dependent divergence in mouse appetitive spatial learning and decision strategies

**DOI:** 10.64898/2026.04.13.718350

**Authors:** Jiaoru Liu, Denise Manahan-Vaughan, Josué Haubrich

**Affiliations:** International Graduate School of Neuroscience, Ruhr-University Bochum, Germany; Ruhr University Bochum, Medical Faculty, Department of Neurophysiology, Bochum, Germany

**Author notes:** **Correspondence:** Josué Haubrich Ruhr University Bochum, Medical Faculty, Department of Neurophysiology, Universitätsstr. 150, MA 4/150, 44780 Bochum, Germany.

**Keywords:** mouse, strain, aging, spatial, appetitive, learning and memory

## Abstract

Animals rely on associative spatial memory to navigate toward previously learned, reward-associated goals. This reward-guided navigation is supported by the hippocampus and its interactions with cortical and subcortical regions: processes which are vital for integrating sensory cues and forming experience-dependent associations. In adulthood, hippocampal-dependent information processing is shaped by aging, reflecting changes in synaptic plasticity and neuromodulatory support. In parallel, mice with distinct genetic backgrounds show systematic differences in behavior, sensory function, and hippocampal plasticity. How mouse strain and age interact to affect spatial appetitive memories has not been well defined. Here, we trained CBA/CaOlaHsd and C57BL/6 mice in early (2–3 months) or late adulthood (7–8 months), in daily training events, to perform a T-maze task with rewards available at a fixed location and decreasing probability in one maze arm. The task consisted of an initial deterministic phase in which a correct response was always rewarded, followed by a probabilistic phase during which reward omissions became increasingly prevalent. We measured correct choices, as well as latencies across training blocks, and combined trial-by-trial metrics with reinforcement learning modeling to assess decision strategies. We observed that CBA/CaOlaHsd mice displayed lower choice latencies than C57BL/6 mice, reached high performance earlier, and maintained better performance when the reward probability decreased. Age was associated with higher latencies and modulated both performance and decision policies in a strain-dependent manner. Moreover, CBA/CaOlaHsd mice displayed higher learning rates from positive outcomes and adopted strategies consistent with more robust context exploitation under uncertainty. C57BL/6 mice, by contrast, exhibited stronger omission-driven strategy switching. Together, these findings reveal robust strain- and age-dependent differences in trial-by-trial decision policies and spatial learning performance.

## Introduction

Animals rely on memory to navigate toward previously learned reward-associated goals (Sosa and Giocomo, 2021). Interactions between the hippocampus and distributed cortical and subcortical regions support reward-guided navigation, allowing sensory cues to be integrated with spatial-contextual representations during memory formation and retrieval (Jiang et al., 2022; Stacho and Manahan-Vaughan, 2022; Haubrich et al., 2025b). In mice, these processes are known to be altered by strain and age (Kim et al., 2008; Wiescholleck et al., 2014; Beckmann et al., 2020; Febo et al., 2024), but the exact impact that they exert on spatial appetitive learning is not clearly defined.

Age is a key biological factor that shapes the formation and updating of spatial and contextual memories across adulthood (Lester et al., 2017). Compared to younger (7-8 week old) animals, moderately aged (7-8 month old) rodents already show deficits in hippocampus-linked cognitive domains, including context-dependent extinction learning (EL) and renewal (Wiescholleck et al., 2014; Zhao et al., 2023), as well as object recognition and object-place learning (Wiescholleck et al., 2014). Mechanistically, age effects on hippocampal-dependent memory are linked to alterations in hippocampal synaptic plasticity and neuromodulation that typically supports its persistence (Foster, 1999). Dopaminergic (DA) and noradrenergic (NA) signaling via D1/D5 receptors (D1/D5R) and beta-adrenergic receptors (beta-AR) are especially important for conferring salience prioritization and promoting experience-dependent long-term hippocampal plasticity (Babushkina and Manahan-Vaughan, 2022; Hagena and Manahan-Vaughan, 2025). Consistent with an age-related weakening of catecholaminergic neuromodulation from the ventral tegmental area and locus coeruleus (Pereira et al., 2021; Langley et al., 2022; Dahl et al., 2023; Shan et al., 2023; Sagheddu et al., 2024), catecholaminergic agonists that facilitate persistent hippocampal plasticity in young adults (7-8 weeks old rats) lose efficacy in moderately aged rats (7-8 months old) (Twarkowski and Manahan-Vaughan, 2016). This may reflect age-related reductions in hippocampal D1/D5 and β-adrenergic receptor binding (Araki et al., 1997; Suzuki et al., 2001) and expression (Popova and Petkov, 1989), less efficient downstream intracellular signaling (Reis et al., 2005), and increased susceptibility to receptor desensitization during agonist stimulation (Bouvier et al., 1988; Hausdorff et al., 1989; Ng et al., 1994; Twarkowski and Manahan-Vaughan, 2016). In addition, moderately aged rats (7-8 months old) show a decline in attention and vigilance, which may further shape the engagement of hippocampal-dependent memory processes (Guidi et al., 2015).

Strikingly, systematic differences among inbred mouse strains have been reported with regard cognition, locomotion, exploration, sociability, empathy, and anxiety-like behavior, all of which can affect performance in learning tests and undermine cross-study comparisons (Kim et al., 2008; Seibenhener and Wooten, 2015; Seemiller et al., 2021; Panksepp and Lahvis, 2023). Direct comparisons show that common laboratory strains, such as C57BL/6 and CBA mice (e.g., CBA/J or CBA/CaOlaHsd), differ in novelty-related behavior, anxiety-like responses, and spatial memory (Sultana et al., 2019; Beckmann et al., 2020) — for example, exploration tests showed higher activity in C57BL/6J than CBA/J (implying stronger novelty-driven exploration), while anxiety-associated measures were higher in CBA/J (implying greater stress responsivity) (Sultana et al., 2019).

In addition, C57BL/6 mice exhibited spatial memory deficits compared to CBA/CaOlaHsd mice, failing to increase exploratory behavior in response to the spatial reconfiguration of familiar objects (Beckmann et al., 2020). Behavioral divergence is not limited to comparisons between C57BL/6 and CBA strains (e.g., CBA/J or CBA/CaOlaHsd) (Kim et al., 2008; Seemiller et al., 2021; Sheppard et al., 2022). Fear conditioning studies have shown that in this task mice exhibit a range of behaviors beyond freezing, such as rearing and grooming, and that these responses differ across inbred strains (including C57BL/6J, DBA/2J, FVB/NJ, SWR/J, BTBR T+ Itpr3tf/J, SM/J, LP/J, and 129S1/SvImJ) (Seemiller et al., 2021). Moreover, C57BL/6 mice display a distinct gait and posture compared to other strains, characterized by a higher amplitude of tail and nose movements (Sheppard et al., 2022).

Sensory function is another major source of divergence across mouse strains: C57BL/6 mice exhibit early onset, progressive hearing loss that typically begins at high frequencies (basal cochlea) by 3 months. They also exhibit congenital ocular dysfunction (Moore et al., 2018). In contrast, CBA strains generally maintain comparatively stable auditory function during early life and show minimal hearing loss until they are 18 months old (Mikaelian, 1979; Walton et al., 1995; Park et al., 2010). They also have healthy retina and ocular function (Feldmann et al., 2019). Sensory decline can drive cortical reorganization and broad changes in plasticity-related receptor expression (including alterations in glutamatergic NMDA and GABAergic receptors) (Feldmann et al., 2019; Beckmann et al., 2020; Hagena et al., 2022a, 2022b), and in C57BL/6 mice is associated with altered hippocampal information processing, impaired synaptic plasticity, and spatial memory deficits, relative to sensorily intact CBA/CaOlaHsd mice (Beckmann et al., 2020; Hagena et al., 2022a). Beyond sensory factors, hippocampal plasticity differs by strain: in freely behaving mice, high-frequency stimulation of the Schaffer collateral–CA1 pathway induces LTP of greater magnitude (lasting >24 h) in CBA/CaOlaHsd than in C57BL/6 mice. This is accompanied by strain-dependent dopaminergic D1/D5 receptor-support of LTP and higher hippocampal D1 receptor expression in CaOlaHsd (Hagena et al., 2022b). Therefore, genetic background in mice should be treated as an important biological variable for learning, rather than as experimental “noise” (Brooks et al., 2004; Sultana et al., 2019; Seemiller et al., 2021; Caliskan et al., 2022; Sheppard et al., 2022; Orozco-Coles et al., 2026). However, a thorough understanding of how genetic background and age shape learning in mice, particularly appetitive and spatial learning, is still lacking.

To address this question, we compared 2-3-month-old and 7–8-month-old mice from two strains (C57BL/6 and CBA/CaOlaHsd) in an appetitively motivated T-maze spatial learning task composed of a deterministic and a probabilistic phase (André and Manahan-Vaughan, 2016; Méndez-Couz et al., 2019). We quantified acquisition performance including correct choices (accuracy) across training blocks and decision time/latency. Collectively, our results reveal specific effects of age, strain, and “age x strain” interactions, which significantly affect task performance, as well as the underlying learning strategies employed by the animals, while they navigate a spatial environment to obtain rewards. These findings highlight the importance of the choice of rodent strain and age, when conducting behavioral assays of cognition.

## Materials And Methods

Experiments were conducted in 2-3-month-old and 7-8-month-old male and female CBA/CaOlaHsd and C57BL/6 mice (Charles River, Germany and Zentrale Versuchstierhaltung der Medizin (ZVM), Ruhr University Bochum). Procedures were performed according to the guidelines of the European Communities Council Directive of September 22nd, 2010 (2010/63/EU) for care of laboratory animals and after prior approval by the local ethics committee (Landesamt für Arbeitsschutz und Ernährung, Nordrhein Westfalen). The mice were housed in temperature- and humidity-controlled vivaria (Scanbur, DK) with a 12-h light-dark cycle (light-period from 6 a.m. to 6 p.m.). All mice had ad libitum access to water. All the mice were pre-handled daily for 5 days and subjected to mild food deprivation before training, and their weights were monitored to maintain 90% of their pre-diet weight.

### Behavioral Apparatus

The experiments were conducted in a T-maze (Fig. 1A) (Haubrich et al., 2025a) composed of a starting box (22cm × 10cm), a center corridor (90 cm × 10 cm), a decision corridor (100 cm × 10 cm), as well as two return corridors (left and right; 85 cm × 10 cm), with 19 cm high walls (Figure 1). The starting box was separated by three sliding doors from the center corridor and from each return corridor. The center corridor connected the starting box to the decision corridor, comprising the left and right T-maze (goal) arms. The two ends of the decision corridor connected to the left and right return corridors, respectively, and each return corridor led back to the starting box. The maze floor and walls were made of gray polyvinyl chloride. Outer visual cues consisted of black-and-white striped 3-dimensional cylinders (50 cm high and 10 cm wide), positioned between the left side corridor and the center corridor, and between the right-side corridor and the center corridor. A white cardboard panel (50 x 50 cm) containing a black circle (25 cm in diameter) was placed on the wall behind the central arm’s end. The reward was always placed on one side at the end of the (e.g. left) goal arm throughout the experiment. and weak odor cues were placed at both end of the goal arms (vanilla scent). The maze was housed in an opaque, dark blue, fabric tent (2.0 m length × 1.85 m width × 2.4 m height) illuminated by white LEDs at 30 lux. The animals’ behavior was monitored using a video camera (Basler GemICam, Basler AG, Ahrensburg, Germany) and Ethovision XT v13 software (Noldus, Wageningen, The Netherlands) for offline analysis. As an appetitive reward, 0.5 cm3 sized food pellets (Dustless Precision Pellets #F05684, Bio-Serv. San Diego, USA) were used.

**Figure 1.**
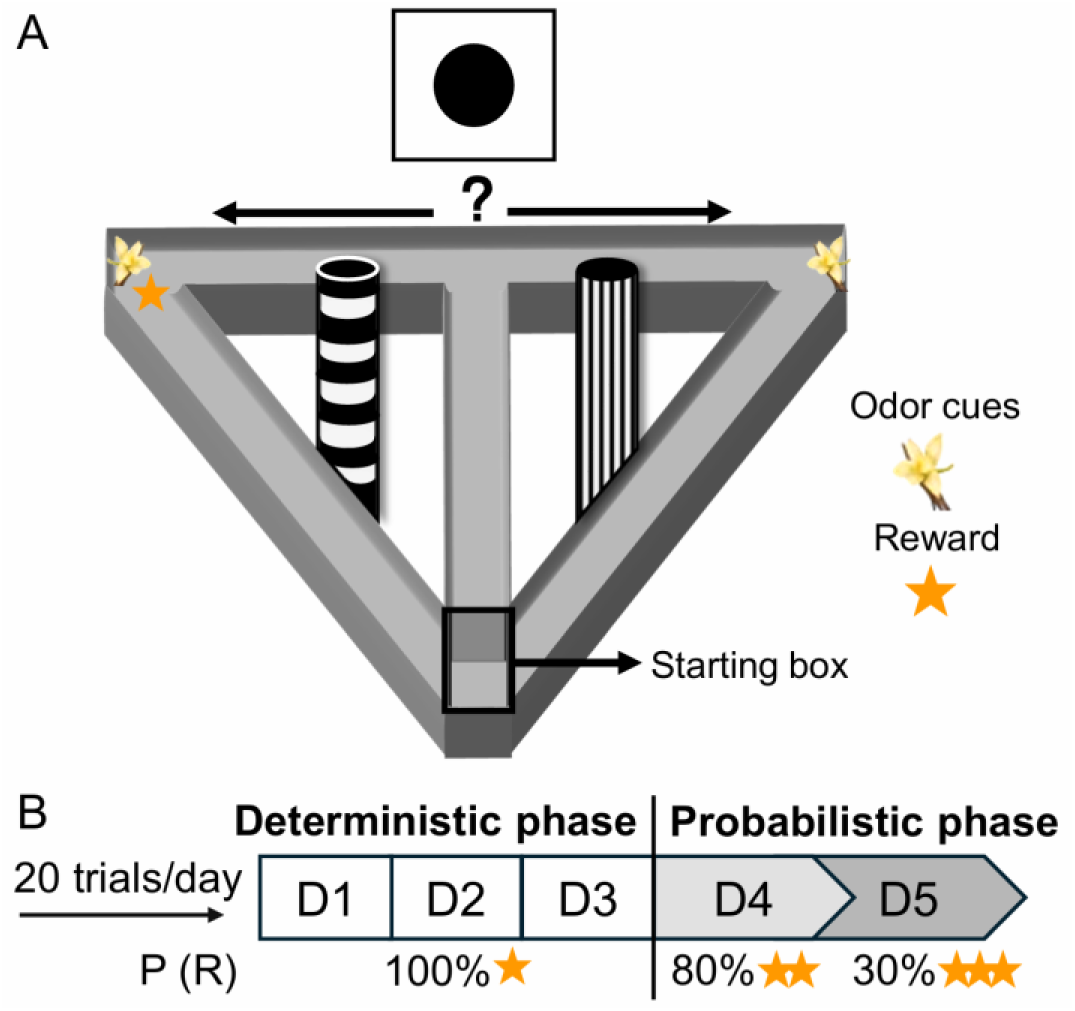
Task design and reward schedule. A) The T-maze was composed of a starting box, central corridor, goal arms, and return corridors. Distal visual cues were positioned around the maze (striped 3D cylinders between the side and center corridors and a cue card with a black circle placed behind the central arm), and a weak odor cue (vanilla) was located at the end of each goal arm. A reward (yellow star) was available in a fixed goal arm (e.g. only left) across trials and days. B) A five-day T-maze protocol with 20 trials/day followed habituation. Days 1-3: deterministic reinforcement (100%) on correct trials. Days 4-5: reward probability, P (R), was progressively reduced from 80% to 30% while reward magnitude increased proportionally.

### Behavioral Procedures

In the week prior to training, animals were habituated to the T-Maze for five days. Throughout the procedure, the maze contained no specific odor or spatial cues. On the first three days of habituation, mice were placed in the starting box, that contained scattered food rewards for 10 min per day, and. the door leading to the other areas of the maze was closed. On the fourth and fifth days of habituation, the door between the starting box and maze was opened, allowing them to explore the maze and locate reward pellets that were scattered throughout the T-maze, thereby encouraging exploration. Once the animals reached a goal zone, the center corridor was blocked to encourage the animals’ return to the starting box via the outer T-maze corridors, thus ending a trial. On the fourth day of habituation, the food reward was scattered in both outer corridors and goal zones, but not in the center corridor. On the fifth day of habituation, the food reward was placed only in the goal zones located at the end of the goal arms on the last two habituation days, and mice explored the T-maze for 3-6 trials each day (until a least one reward pellet was consumed). During the last two days of habituation, choice preferences were recorded, and the correct goal zone for each mouse was assigned to the side opposite its preferred T-maze arm.

In the week following habituation, the mice underwent five days of training, each consisting of four blocks of five trials of maximally 2 min duration each. Each trial was separated by an inter-trial interval of 15 s. One arm of the T-maze served as the goal zone (e.g., a right turn into a designated arm) and was associated with reward delivery. Entries into this rewarded location were counted as correct choices, whereas entries into the opposite, non-rewarded arm (the error zone) were counted as incorrect choices. The goal and error zones were kept constant across trials. When a mouse entered either the correct or the incorrect goal zone, its return to the center corridor and the opposite T-maze arm was immediately blocked by a plastic barrier, ensuring that each trial provided only a single choice opportunity. The mouse was then moved back to the starting box via the outer corridor, prior to commencing the next trial. If an animal failed to reach the end of either goal arm within 2 min, it was gently guided back to the starting box, and the trial was ended; these trials were also scored as incorrect choices.

Training comprised two phases: a deterministic phase and a probabilistic phase (Fig. 1B). To promote stable learning of the correct choice, reward delivery was initially deterministic (100% probability) on Days 1–3. To examine how changes in reward probability influenced choice performance and decision updating, reward probability was progressively reduced: to 80–60% on Day 4, to 60–50% across the first 10 trials of Day 5, and to 30% across the last 10 trials of Day 5. To maintain motivation, the number of pellets delivered on some rewarded trials was increased proportionally, in line with the decrease of reward probability, ensuring that in any session, the animals were allowed to obtain up to 20 pellets.

### Analysis of Trial-by-trial Strategies

For each trial transition, the previous trial outcome was classified as a ‘win’ when a reward pellet was obtained, and as a loss if the animal failed to make a correct goal arm choice. ‘Stay’ and ‘shift’ responses were defined as repeating the same arm choice in two consecutive trials versus switching to the opposite arm from one trial to the next, respectively. Conditional probabilities were then computed within each day (20 trials/day) for win-stay, win-shift, lose-stay, and lose-shift. In some cases, arm choices were additionally averaged across days to obtain summary measures for deterministic phases (days 1-3) and probabilistic phases (days 4-5) for each individual animal. During the probabilistic phase, we also computed the probability of staying versus shifting following a correct choice, either collapsing across rewarded or unrewarded correct trials, or stratifying transitions based on whether the preceding correct trial was rewarded or unrewarded. To assess reward magnitude effects, shifts following correct arm choices and rewarded trials were further stratified by the number of pellets obtained (1-4) and analyzed across days 4-5.

### Reinforcement-learning model

To gain better insight about how animals update their choices, based on reward presence and reward omission feedback, we fit a reinforcement-learning (RL) model to each animal’s trial-by-trial behavior. The model tracks the expected reward (Q) for choosing each option (Right/correct arm vs Left/incorrect arm), and values were initialized at q0 = 0.5 for both arms. The model uses a Rescorla-Wagner update rule based on prediction errors (Rescorla RA and Wagner AR, 1972), and the Q values of each arm of the maze were updated separately, consistent with Q-learning models (Watkins and Dayan, 1992; Clifton and Laber, 2020). After each trial t, only the chosen arm was updated using a prediction error, calculated as:

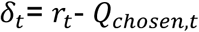

where *r****_t_*** is the obtained reward magnitude (number of pellets). The Q value of the chosen arm is then updated as follows:

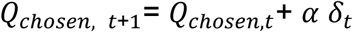

A central feature of this model is that it uses separate learning rates for positive (referred here as α+ or α-pos) and negative outcomes (α- or α-neg). Both learning rates were fit per animal and constrained to the interval (0, 1):

**-** α+ when δ***_t_*** ≥ 0 (better-than-expected outcome; reward) α- when δ***_t_*** < 0 (worse-than-expected outcome; reward omission)

To connect these learned values to choices, the model converts the value difference between arms (Q***_R,t_*** – Q***_L,t_***) into a probability of choosing the correct/right arm. Because the task has two options, we used a logistic (softmax) choice rule(Wilson and Collins, 2019), which maps any value difference to a probability between 0 and 1:

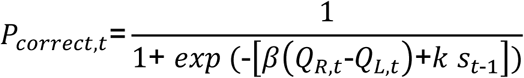

where:

- (Q***_R,t_*** – Q***_L,t_***), or ΔQ, indicates the model’s current preference for the correct/right arm (larger values favor the right arm)
- β is the “inverse temperature”, controlling how strongly choices follow the value difference (larger values favor deterministic choices, lower values favor random responding)
- k captures a tendency to repeat the previous choice, independent of learning (e.g. “stickiness” or perseveration).
- ***s _t-1_*** encodes the previous choice as +1 for right/correct and -1 for left/incorrect (set to 0 on the first trial)

Finally, to allow occasional random choices (e.g., lapses of attention or motor errors), the model included a lapse rate epsilon, which is the probability that the animal chooses randomly (50/50) on a given trial, instead of following learned values:

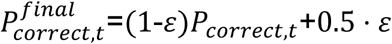

#### Model Fitting

Model parameters were fit by minimizing the negative log-likelihood (NLL) of the observed choices (Wilson and Collins, 2019; Pupillo et al., 2023). The NLL measures how well the model predicts the animal’s choices: if the model assigns high probability to the choices the animal actually made, NLL is low; if it assigns low probability, NLL is high.

Learning rates (α+, α−) and stickiness (k) were fit per animal. In contrast, β and ε were treated as global hyperparameters shared across animals and selected by grid search. We evaluated all (β, ε) pairs in a predefined grid and chose the combination with the best overall fit (Pupillo et al., 2023), using a convergence-aware criterion that penalized solutions with non-converged animals. Optimization used L-BFGS-B with bounds: α+ and α− in (1e−6, 1−1e−6); k in [−5, +5]. Each fit used multi-start optimization (multiple initial parameter seeds) and the best solution (lowest NLL) was retained. Optimizer settings used up to 4000 iterations per fit, with an optional “rescue” step in which non-converged animals were re-fit using larger iterations and a wider set of starting points.

For each animal, the fit yielded estimates of α+, α−, and k, along with trial-wise predicted choice probabilities p ***_correct(t)_*** and the latent value trajectories (Q***_R,t_*** – Q***_L,t_***, and Q***_R,t_*** −Q***_L,t_***). These outputs were used for group comparisons and for visualizing dynamics across training.

#### Model Validation

To verify that the fitted learning rates reflect true latent learning parameters, we performed parameter recovery using simulated datasets. Synthetic data were generated with the same trial-by-trial reward schedule as the real task (including reward probability and magnitude changes across days). For each simulated animal, “true” values of α+, α− and k were sampled from predefined ranges based on the observed experimental data(Wilson and Collins, 2019), and choices were generated by the same RL update and choice equations described above. The same fitting pipeline was then applied to these synthetic datasets. Recovery was quantified by comparing fitted vs true parameters using Pearson and Spearman correlation analyses to confirm that the pipeline can recover the target latent variables.

## 1. Statistics

Following conformation that data full filled a normal distribution, group differences were assessed using two-way factorial analysis of variance (fANOVA) to test for main effects of strain and age, and their interaction, or two-way repeated-measures ANOVA (rANOVA) to examine these effects across testing sessions. When appropriate, Tukey’s post hoc tests were applied to determine specific pairwise comparisons. Correlational analyses were performed using both Pearson’s and Spearman’s correlation tests. Statistical significance was set at α = 0.05 for all tests.

All analyses and plots were generated using R version 4.3.0 (R Foundation for Statistical Computing, Vienna, Austria).

## Results

### Murine spatial learning under deterministic and probabilistic reward schedules reveals strain- and age-dependent differences

We assessed spatial appetitive learning in young (2–3-month-old) and older (7–8-month-old) adult C57BL/6 (Younger C57BL/6, n=16; Older C57BL/6, n=12) and CBA/CaOlaHsd (Younger CBA/CaOlaHsd, n=14; Older CBA/CaOlaHsd, n=12) mice using a five-day T-maze protocol (20 trials/day; Figure 1). During Days 1-3, correct choices were always reinforced in a fixed goal arm, whereas during Days 4-5, reward probability was gradually reduced to 30% while reward magnitude increased proportionally (Figure 1B).

Performance in both mouse strains, expressed as the percentage of correct choices in 5-trial blocks, increased across training (main effect of Block: repeated-measures ANOVA, F_(11, 572)_ = 29.12, p < 0.001; Figure 2). There was no main effect of age (p > 0.05), but there was a main effect of strain (F_(1, 50)_ = 11.21, p = 0.002), with CBA/CaOlaHsd mice (n=26) performing significantly better than C57BL/6 mice (n=28) (Figure 2A-B). This strain difference depended on training stage, as indicated by a significant strain x block interaction (F_(11, 572)_ = 1.84, p = 0.042). Tukey post hoc tests revealed that CBA/CaOlaHsd mice already outperformed C57BL/6 mice by the end of Day 1 (blocks 3-4; p < 0.05) and maintained higher performance during early to mid-training (blocks 7 and 11; p < 0.05). When reward probability was reduced during the final blocks, correct choices declined in C57BL/6 mice (blocks 19–20; p < 0.05), whereas CBA/CaOlaHsd mice maintained stable performance. We also detected a significant Strain x Age x Block interaction (F_(6, 320)_ = 2.84, p = 0.009; Figure 2C). Post hoc analyses indicated that young C57BL/6 mice performed worse than all other groups in the final block (p < 0.001).

**Figure 2.**
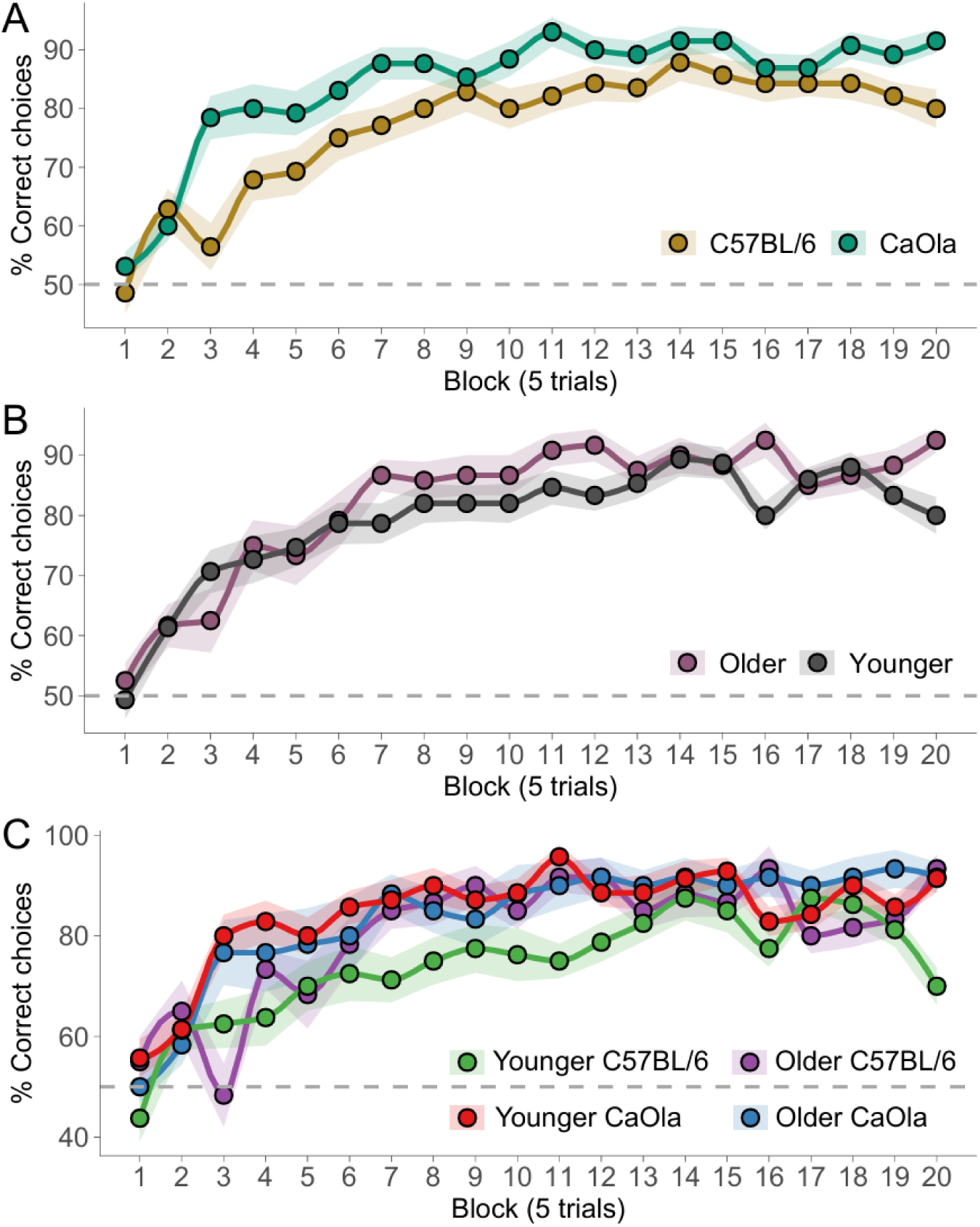
Assessment of spatial learning performance reveals strain- and age-dependent effects across deterministic and probabilistic phases. Spatial learning performance expressed as percentages of correct choices across blocks of 5 trials. The deterministic phase spanned blocks 1 to 16, and the probabilistic phase spanned blocks 17 to 20. Across training, all groups improved performance, but CBA/CaOlaHsd mice reached higher levels of correct performance earlier than C57BL/6 mice and maintained this advantage when reward probability decreased during the probabilistic phase. A) Performance stratified by Strain. B) Performance stratified by Age. C) Performance stratified by Strain x Age group. Points represent group means, and shaded ribbons indicate the standard error of the means

Together, these data indicate a robust strain-dependent difference in appetitive spatial learning: CBA/CaOlaHsd mice acquired the rewarded-arm preference more rapidly and preserved performance when reward delivery became probabilistic. In contrast, C57BL/6 mice showed delayed acquisition and a selective deficit in maintaining learned responses under low reward probability. This effect was especially evident in younger C57BL/6 mice.

### Age and strain-dependent differences in goal-reaching latency are evident

To assess changes in task execution speed, we analyzed the animal’s latencies to reach a goal zone (Figure 3). Latencies decreased across training (rANOVA, main effect of Block: F_(6, 320)_ = 14.24, p < 0.001). When goal arm latencies were assessed across trials and pooled in blocks of 5 trials, a significant age x block interaction was observed (F_(6, 320)_ = 3.15, p = 0.004; Figure 3B), driven by longer latencies in older animals during early blocks (p < 0.05). We also detected a significant Strain x Block interaction (F_(6, 320)_ = 2.84, p = 0.009; Figure 3B), with CBA/CaOlaHsd mice showing lower latencies than C57BL/6 mice across multiple blocks (all trial blocks of day 2, and first and last trial blocks of day 3; Tukey post hoc, p < 0.05), whereas C57BL/6 were faster in block 19 (i.e. the third trial block of day 5) (p < 0.01).

**Figure 3.**
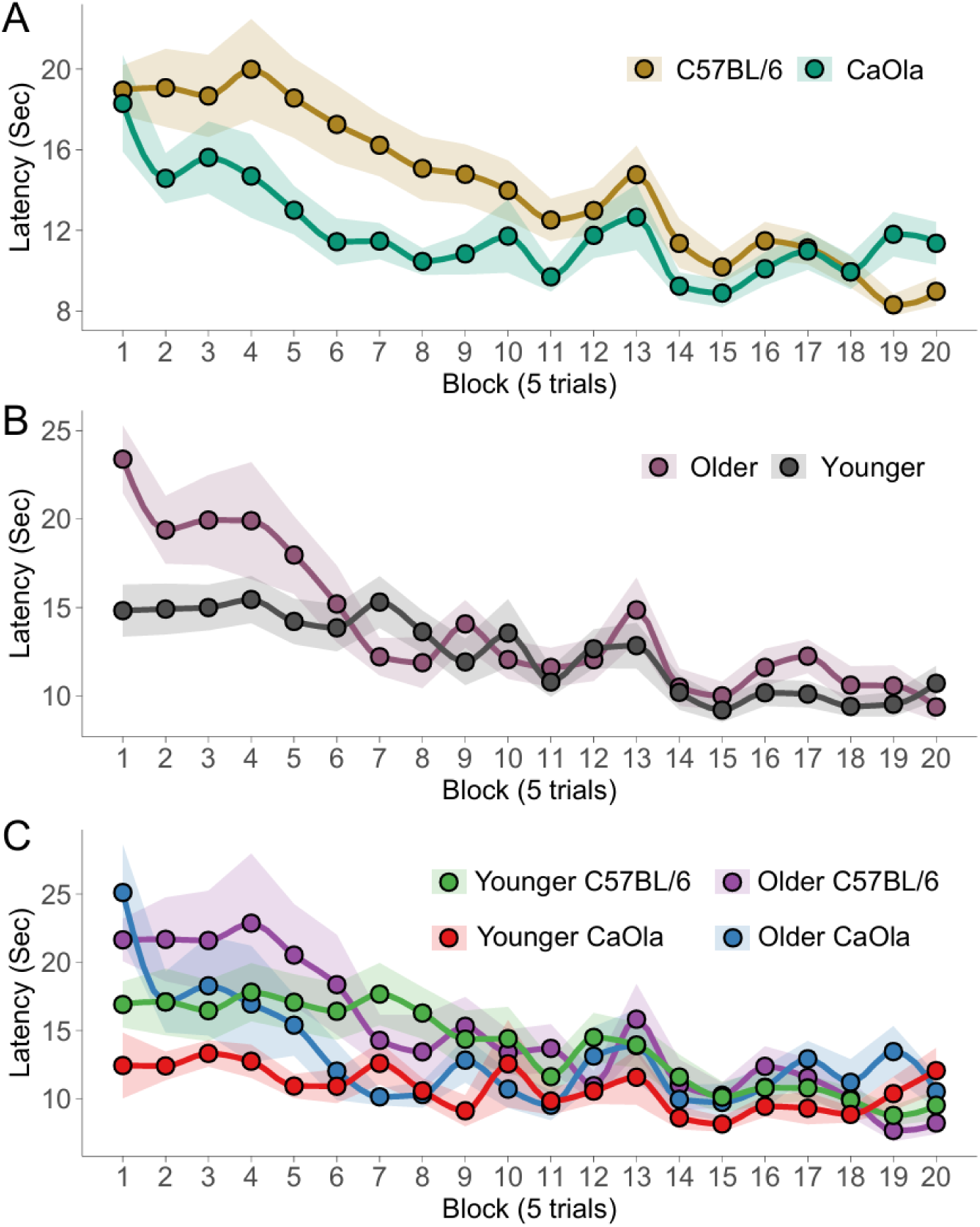
Assessment of goal-reaching latency during spatial learning reveals strain and age-dependent effects that are prominent during the deterministic learning phase. Mean latency (s) to reach a goal zone per 5-trial block. Latencies decreased across training in all groups, consistent with learning. Younger animals were overall faster than older animals, and CBA/CaOlaHsd mice exhibited shorter latencies than C57BL/6 mice during early and mid-deterministic phases (blocks 1-16), indicating more efficient task acquisition. Differences were attenuated during the probabilistic phase (blocks 17-20) A) Latency stratified by Strain. B) Latency stratified by Age. C) Latency stratified by Strain x Age group. Points represent group means, and shaded ribbons indicate the standard error of the means.

Thus, as expected, younger animals were faster overall in reaching the goal zones. Notably, as animals learned to navigate the maze to obtain rewards, their latencies progressively decreased, with this improvement emerging earlier in CBA/CaOlaHsd mice.

### Strain and age-dependent divergence is apparent in trial-by-trial decision patterns

To quantify trial-by-trial decision strategies, we computed the probability of repeating (stay) or switching (shift) the previous choice, for each individual mouse, conditional on whether the preceding trial was rewarded (win) or unrewarded (lose) (Figure 4). These indices were calculated within each day (20 trials/day).

**Figure 4.**
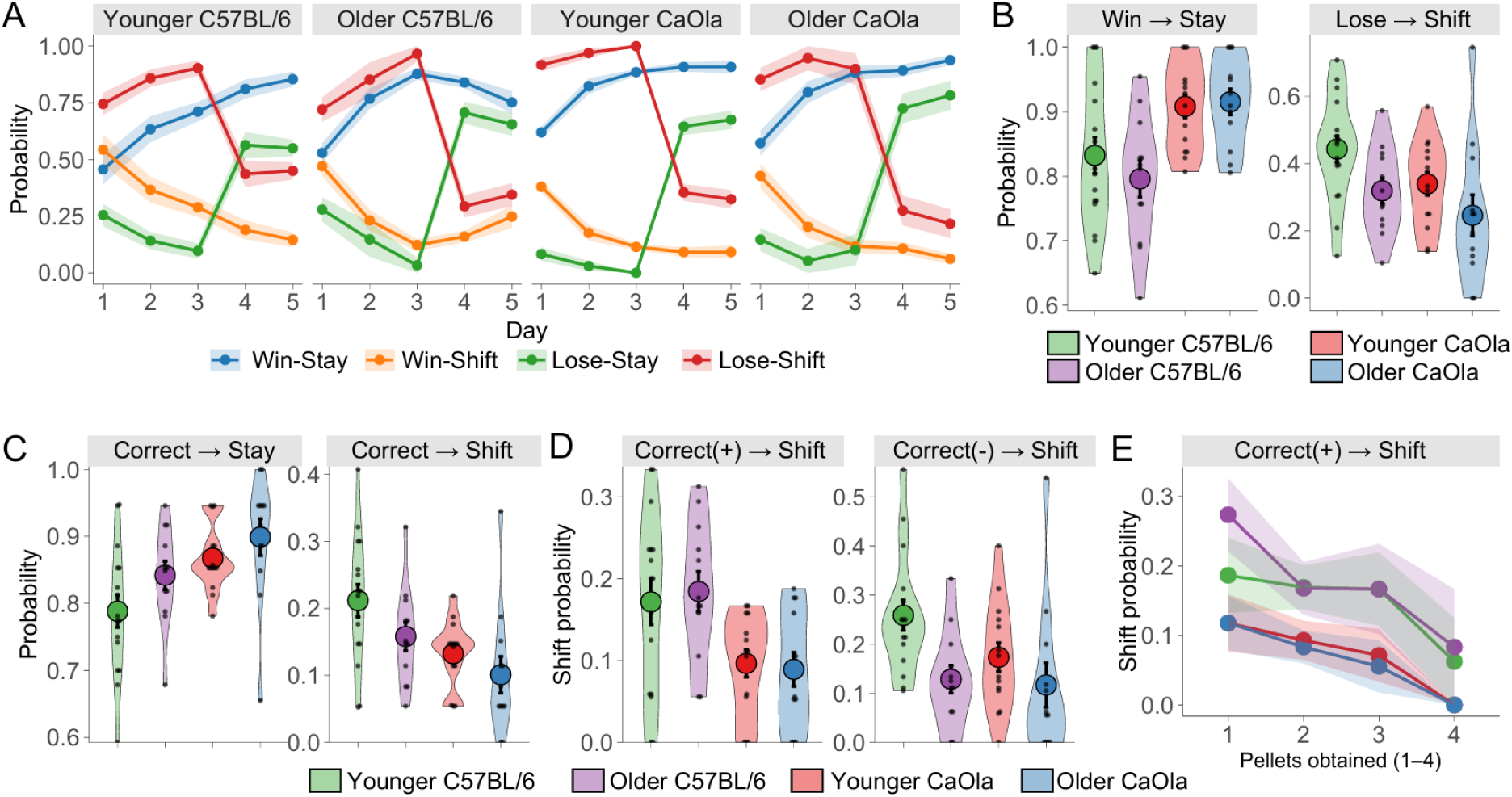
Strain and age shape trial-by-trial decision strategies and perseverance under uncertainty. Trial-by-trial analyses reveal how animals adjusted their choices following rewarded and unrewarded outcomes and how these strategies evolved across deterministic (days 1 to 3) and probabilistic phases (days 4 and 5). CBA/CaOlaHsd mice adopted adaptive win-stay/lose-shift strategies earlier than C57BL/6 mice and showed greater perseverance when reward delivery became probabilistic. Younger animals were more sensitive to reward omission, whereas older animals exhibited greater stability of learned choices in the probabilistic phase. A) Probabilities of Stay or Shift responses following Win or Lose outcomes across Days 1-5. During the deterministic phase, CBA/CaOlaHsd mice showed earlier differentiation between adaptive and non-adaptive strategies. By Day 3, all groups displayed clear win-stay and lose-shift asymmetries. B) Average Win/Stay and Lose/Shift during the probabilistic phase (Days 4 -5). CBA/CaOlaHsd mice maintained stronger adaptive responses under uncertainty. C) Average probability of Stay or Shift responses following a correct choice during the probabilistic phase. CBA/CaOlaHsd mice showed higher correct-stay behavior. D) Shift probability following a correct rewarded trial versus a correct unrewarded trial during the probabilistic phase. C57BL/6 mice showed higher shift probabilities, and younger animals showed higher shifting specifically after losses. E) Shift probability following a correct rewarded trial during the probabilistic phase, stratified by reward magnitude. Strain differences were reduced at the highest reward magnitude. For panels A and E, points indicate group means and shaded ribbons indicate SEM. For panels B–D, large points show group means, error bars indicate SEM, and small points show individual animals.

We found that the outcome-dependent choice strategy evolved across the task (Figure 4A). For this we assessed whether prior experience impacted on the next arm choice. This. allowed us to gain insights into behavioral characteristics such as perseverance, resilience and adaptability. This was demonstrated by the finding that within each Age x Strain group, a repeated-measures ANOVA with the factors Metric (win-stay, win-shift, lose-stay, lose-shift) and Day (1-5) revealed a strong main effect of Metric and a Metric x Day interaction in all groups (all p < 0.001). We also detected a significant Strain x Age x Day interaction for metrices (F_(2.53, 126)_ = 3.57, p = 0.02), revealing that strain differences, in the evolution of outcome-dependent strategies, varied as a function of age. In the initial deterministic phase, post hoc comparisons showed that CBA/CaOlaHsd mice exhibited pronounced differentiation in decision strategies following wins and losses already on Day 1, including a clear separation between lose-stay and lose-shift (p < 0.001) and win-stay and win-shift (p > 0.001), indicating that their choices were already in a direction that maximized reward acquisition. In contrast, although C57BL/6 mice also display a robust asymmetry between lose-stay and lose-shift (p < 0.001), they showed no detectable difference between win-stay and win-shift on Day 1 (p = 0.17). By Day 3, the contrast between win-stay and lose-shift remained detectable in younger groups (younger C57BL/6: p < 0.005; younger CBA/CaOlaHsd : p < 0.001) but not in older animals (older C57BL/6: p = 0.148; older CBA/CaOlaHsd: p = 0.995), indicating that younger mice were more strongly driven by loss-related feedback, whereas older animals displayed a more balanced reliance on win- and loss-related feedback. Importantly, by Day 3, all groups exhibited strong asymmetries between lose-stay and lose-shift and between win-stay and win-shift (all p < 0.001).

By Day 5, in the probabilistic phase, loss-contingent indices continued to diverge in CBA/CaOlaHsd and in older C57BL/6 (all p < 0.001), whereas younger C57BL/6 no longer showed a significant difference between lose-stay and lose-shift (p = 0.193). In parallel, win-stay and lose-stay became indistinguishable in older animals (older C57BL/6: p = 0.52; older CBA/CaOlaHsd: p = 0.11) but remained distinct in both younger groups (all p < 0.001), indicating that older animals relied more strongly on the previously acquired task rule during the probabilistic phase, whereas younger mice remained more sensitive to immediate feedback.

Because reward contingencies changed on Days 4-5 (decreased reward probability with increased reward size), we next focused on this probabilistic phase (Figure 4B). After averaging each metric across Days 4-5 within animals, fANOVA of win-stay behavior revealed a significant main effect of Strain, with CBA/CaOlaHsd showing higher win-stay probabilities than C57BL/6 mice (F_(1,50)_ = 14.8, p < 0.001), with no effect of Age and no Strain x Age interaction (p > 0.05). In contrast, lose-shift behavior depended on both Strain (F_(1,50)_ = 4.34, p = 0.042) and Age (F_(1,50)_ = 6.33, p = 0.015), with no Strain x Age interaction (p = 0.7): lose-shift was lower in CBA/CaOlaHsd than C57BL/6 mice (p = 0.035) and lower in older than younger mice (p = 0.015), indicating greater persistence in maintaining the learned strategy despite occasional non-reward following correct choices.

We next examined decision updating during the probabilistic phase, specifically after correct arm choices. This was done either by collapsing stay and shift responses across reward delivery outcomes (Figure 4C), or stratifying trials into those in which reward was obtained, or not (Figure 4D). When collapsing across reward delivery, CBA/CaOlaHsd mice showed higher correct-stay and correspondingly lower correct-shift behavior than C57BL/6 mice (fANOVA: F_(1,50)_ = 9.81, p = 0.003 for both measures), indicating that CBA/CaOlaHsd displayed greater persistence in repeating correct choices despite uncertainty in reward delivery. Older animals showed a non-significant trend toward higher correct-stay and lower correct-shift (p = 0.06), with no evidence for a Strain x Age interaction (p = 0.62), also suggesting a tendency toward greater persistence in older animals.

When we stratified correct-shift behavior by reward outcome, shifts following a correct and rewarded trial differed between strains, with C57BL/6 mice showing higher shift probabilities (fANOVA: F_(1,50)_ = 12.8, p < 0.001), indicating a tendency to abandon previously rewarded choices. In contrast, shifts following a correct but unrewarded trial were higher in younger than older animals (F_(1,50)_ = 7.77, p < 0.05) with no main effect of strain or age x strain interaction (p > 0.05), indicating stronger behavioral updating after reward omission in younger mice. Finally, stratifying shifts after correct and rewarded trials by reward magnitude (Figure 4E) showed that strain differences were present for small-to-intermediate rewards (1-3 pellets; p < 0.05) but not for the largest reward (4 pellets; p = 0.8), suggesting that high reward magnitude reduced exploratory switching and attenuated strain-dependent differences in shift strategies.

Therefore, CBA/CaOlaHsd mice displayed better initial learning of task rules and higher perseverance during the probabilistic phase relative to C57BL/6 mice. Age primarily influenced responsiveness to reward omission, with younger animals showing greater shifting after unrewarded correct trials, suggesting reliance on immediate feedback and working-memory, whereas older animals were more likely to rely on the learned rule. Notably, the highest reward magnitude reduced strain differences in post-win shifting, indicating that strong reinforcement stabilized this behavior across groups.

### Reinforcement learning explains strain-dependent differences in decision updating

The strain-dependent differences in trial-by-trial decision strategies that we detected in C57Bl/6 and CBA/CaOlaHsd mice suggest that animals may differ in how they learn from outcomes and that this learning characteristic is strain-dependent. In the behavioral task used here, mice can update their choice policy based on rewarding outcomes (the correct arm yields a reward) and unrewarded outcomes (reward omission or choosing the incorrect arm). With training, animals typically overcome an innate alternation tendency and develop a stable preference for the baited arm. Therefore, we hypothesized that group differences in learning progression and decision-making could be explained by differences in learning rates related to positive versus negative outcomes. To test this, we fit a two-learning-rate Rescorla-Wagner/Q-learning model (Cazé and Van Der Meer, 2013; Lefebvre et al., 2017; Kastner et al., 2020) to the trial-by-trial choices of each individual animal. In our model, the value of selecting an arm is updated separately following rewarded outcomes (learning rate α-pos) versus unrewarded outcomes, including reward omissions during correct choices (learning rate α-neg). In addition, the model included an individual stickiness parameter (k) to capture each mouse’s outcome-independent tendency to repeat the previous choice (perseveration; positive k) or alternation of arm choice across trials (negative k).

We first assessed the model’s ability to recover the targeted latent variables. This served to determine whether the model can accurately recover the underlying learning parameters from choice behavior. We generated synthetic datasets using the same task structure, simulating agents with known α-pos, α-neg, and k values, and then refit the model to these simulated choices. The model successfully recovered these parameters, as indicated by strong correlations between true and estimated values for α-pos (Pearson p < 0.01; Spearman p < 0.0001), α-neg (Pearson p < 0001; Spearman p < 0.0001), and k (Pearson p < 0005; Spearman p < 0.001). After confirming parameter recoverability, we fit the model to the empirical data. Model-predicted choice probabilities tracked the observed choice behavior across groups (Figure 5A), with a strong overall association between predicted and observed responses (Pearson r = 0.88, p < 0.0001), and similarly strong associations within each group (all r ≥ 0.8, all p < 0.001).

**Figure 5.**
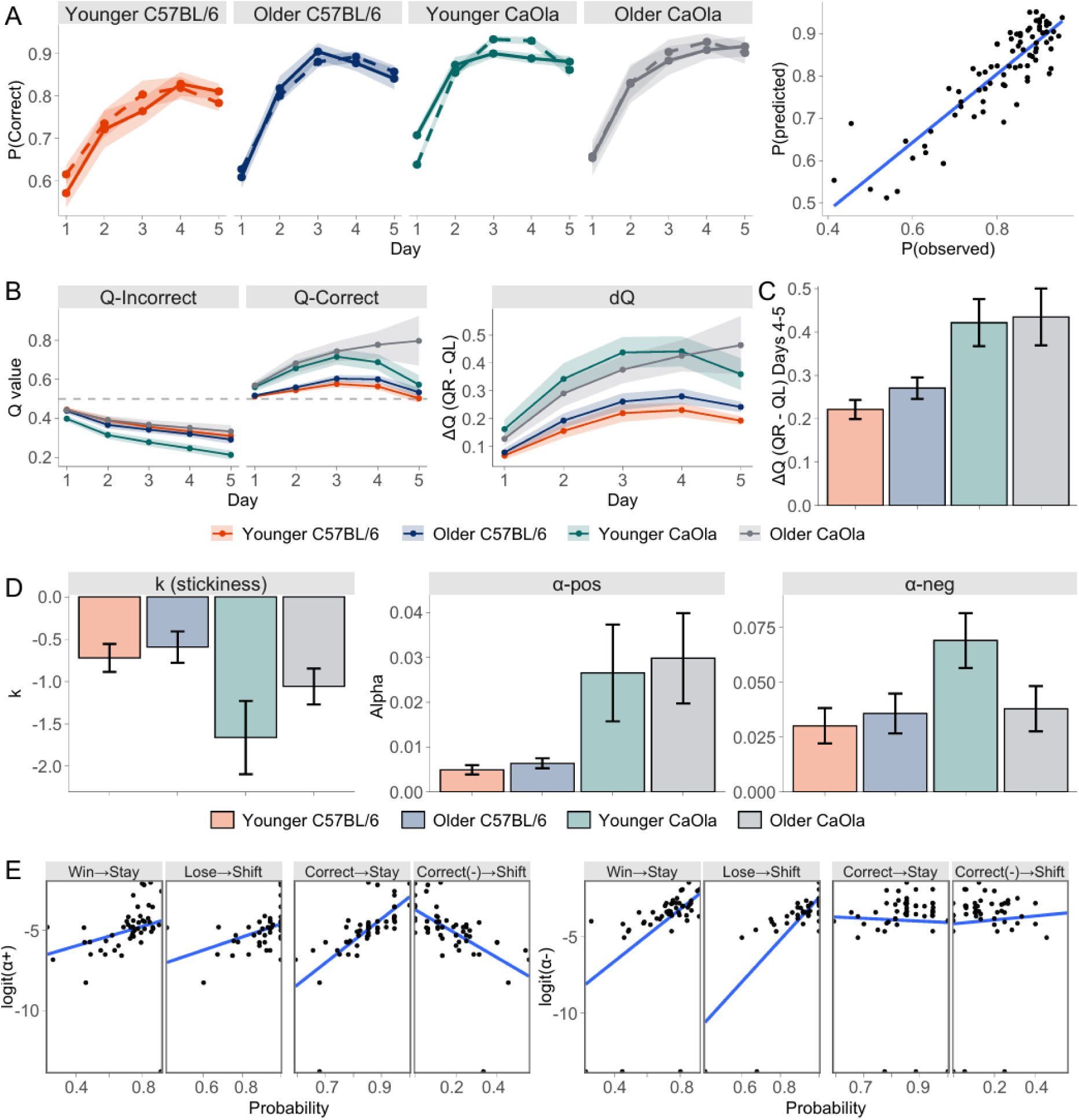
Reinforcement learning modeling reveals stronger reward-driven updating and more stable value representations in CBA/CaOlaHsd mice. Computational modeling shows that strain differences in spatial learning are driven primarily by enhanced updating following rewarded outcomes and greater stability of learned values in CBA/CaOlaHsd mice. A) Model validation. Left: observed probabilities of correct choices (solid lines; ribbons indicate ± SEM) overlaid with model-predicted probabilities (dashed lines). Right: scatterplot of observed versus predicted probabilities of correct choices, with the blue line indicating the line of best fit. B) Q values across training. Lines show group means and ribbons indicate SEM for Q values of the incorrect arm (left), the correct arm (middle), and their separation (ΔQ; right). CBA/CaOlaHsd mice developed a larger value difference between arms, reflecting a stronger and more stable preference for the correct option. C) ΔQ during the probabilistic phase. Bars show group means and error bars indicate SEM for ΔQ during the probabilistic phase. CBA/CaOlaHsd mice maintained greater value separation under uncertainty, indicating persistence of the learned rule. D) Differences in fitted model parameters. Bars show group means and error bars indicate SEM for stickiness (left), α-pos (center), and α-neg (right). CBA/CaOlaHsd mice exhibited higher α-pos, indicating stronger updating following rewarded outcomes, and lower k, indicating higher outcome-independent tendency to alternate. Moreover, younger CBA/CaOlaHsd additionally showed elevated α-neg, indicating stronger updating following losses. E) Scatterplots showing the relationship between observed Win/Lose-Stay-Shift probabilities and logit-transformed α-pos (left) and α-neg (right), with the blue lines indicating best-fit regressions, revealing reinforcement learning parameters that predict adaptive decision policies.

In the model, Q values reflect the learned expected payoff of each arm (initialized at 0.5). Across groups, Q values decreased for the incorrect arm (Figure 5B, left) and increased for the correct arm (Figure 5B, middle). For the incorrect-arm Q values, a significant Strain x Age interaction was detected (rANOVA: F_(1,46)_ = 6.90, p = 0.01), and Tukey’s post hoc tests indicated that younger CBA/CaOlaHsd mice reduced the expected value for the incorrect arm more strongly than the other groups (p < 0.001), indicating a more pronounced devaluation of the expected reward following negative outcomes. For correct-arm Q values, we observed a significant main effect of strain, with CBA/CaOlaHsd showing higher values than C57BL/6 mice (F_(1,46)_ = 12.3, p = 0.001), suggesting a higher accumulated expected value for the correct arm in CBA/CaOlaHsd mice. Consistent with this, the value separation between arms (ΔQ; Figure 5B, right) was larger in CBA/CaOlaHsd than in C57BL/6 mice (F_(1,46)_ = 14.99, p < 0.001). Restricting the analysis to the probabilistic phase (Days 4-5), when reward probability was low, ΔQ remained higher in CBA/CaOlaHsd (F_(1,46)_ = 15.49, p < 0.001), with no main effect of age and no Age x Strain interaction (both p > 0.5), showing that CBA/CaOlaHsd mice maintained stronger value contrast between the correct and incorrect arms even under uncertainty. Together, these results suggest that CBA/CaOlaHsd formed a stronger value distinction between the correct and incorrect arms across training, driven primarily by a higher correct-arm value (and, in younger CBA/CaOlaHsd, also by a lower incorrect-arm value). In other words, CBA/CaOlaHsd mice developed a stronger and more stable internal representation of which arm was advantageous, mainly because rewarded outcomes strengthened their preference for the correct arm more effectively.

We next examined group differences in the three key fitted parameters: k, α-pos, and α-neg. Stickiness (k) values were more negative in CBA/CaOlaHsd than in C57BL/6 (fANOVA, main effect of strain: F_(1,46)_ = 6.43, p = 0.015), with no main effect of age and no Age x Strain interaction (both p > 0.1) (Figure 5D, left). Negative k indicates a greater outcome-independent tendency to alternate, indicating that CBA/CaOlaHsd mice displayed a higher intrinsic tendency to switch between arms, independent of learning rates.

Next, we compared learning rates for rewarded outcomes (α-pos) and unrewarded outcomes (α-neg). This can reveal whether observed group differences in task performance reflect differences in how strongly animals update their predictions following rewarded versus unrewarded outcomes. Given that learning rates are bounded (0-1) and right-skewed, statistical analyses followed previous reports (Seiffert et al., 2024) and were performed on logit-transformed α values (untransformed values are shown in the figure). fANOVA revealed that α-pos was higher in CBA/CaOlaHsd (F_(1,46)_ = 11.35, p = 0.002; Figure 5D, middle), indicating enhanced learning from positive reinforcement in CBA/CaOlaHsd mice. For α-neg, we detected a significant Age x Strain interaction (F_(1,46)_ = 4.95, p = 0.03), with post hoc tests indicating that younger CBA/CaOlaHsd had higher α-neg than younger C57BL/6 (p = 0.03) (Figure 5D, right). This suggests that younger CBA/CaOlaHsd mice also showed enhanced updating in response to negative outcomes compared to younger C57BL/6 mice. Thus, CBA/CaOlaHsd mice showed stronger learning from rewarded experience overall, and younger CBA/CaOlaHsd additionally showed stronger updating of the arm choice strategies, based on negative outcomes, compared to younger C57BL/6.

Finally, we tested whether these parameters correlated with specific win/lose-stay/shift policies. This was done to determine whether variability in reinforcement learning parameters translated into distinct win/lose-stay/shift policies. During the deterministic phase, logit(α-pos) (Figure 5E, left) correlated positively with Win-Stay (Spearman ρ = 0.50, p < 0.0001) and with Lose-Shift responses (ρ = 0.50, p = 0.001), indicating that α-pos captured the extent to which animals implemented reward-driven adaptive strategies. During the probabilistic phase, logit (α-pos) was strongly positively correlated with staying after a correct choice (ρ = 0.81, p < 0.0001) and negatively correlated with shifting after a correct but unrewarded choice (ρ = -0.70, p < 0.0001). In other words, effective learning derived from memory of the (occasionally) rewarded arm was associated with greater tolerance of omissions in a stable but uncertain environment. This kind of arm choice strategy will depend on confidence in reference memory. Logit (α-neg) (Figure 5E, right) also correlated with Win-Stay (ρ = 0.63, p < 0.0001) and Lose-Shift (ρ = 0.67, p = 0.001) during the deterministic phase, but did not significantly relate to probabilistic-phase measures (all p > 0.1), raising the possibility that decisions during the probabilistic phase may rely less on learned policies and depend more strongly on working memory processes. Stickiness (k) was not associated with these measures (all p > 0.1), except for Lose-Shift during the deterministic phase (ρ = −0.72, p < 0.0001).

Taken together, these findings show that the two strains differ in how they learn and maintain task rules. CBA/CaOlaHsd mice were more strongly influenced by rewarding outcomes (higher α-pos), leading to a clearer and more stable preference for the correct arm (higher ΔQ), even during uncertainty. Importantly, this stability did not reflect simple habitual repetition of previous choices, as these mice also showed a higher general tendency to alternate independently of outcomes (lower k), suggesting instead a more reliable reference memory of which option was advantageous. In contrast, the learning behavior of C57BL/6 mice was comparatively influenced less by reward, resulting in a weaker preference for the correct arm, likely making their behavior more sensitive to recent outcomes.

## Discussion

Rodents display distinct strain-dependent learning (Brooks et al., 2004; Sultana et al., 2019) and hippocampal plasticity profiles (Beckmann et al. 2020; Hagena et al. 2022a; Hagena et al. 2022b), which can be further modulated by age (Wiescholleck et al., 2014; Guidi et al., 2015; Twarkowski and Manahan-Vaughan, 2016; Lester et al., 2017). Consistent with this view, we demonstrate here that C57BL/6 and CBA/CaOlaHsd mice of different ages differ markedly in multiple aspects of spatial appetitive learning. These differences were evident in overall performance and goal-reaching latency, in the evolution of decision strategies, and in measures of reinforcement-driven updating. Specifically, CBA/CaOlaHsd mice acquired task contingencies more rapidly, developed more stable choice preferences under uncertainty, and showed stronger learning from rewarded outcomes, whereas C57BL/6 mice were more sensitive to trial-by-trial outcome fluctuations. Age further shaped these patterns by modulating perseverance and sensitivity to reward omission. Together, these findings reveal age- and strain-dependent differences in how spatial reinforcing associations are formed, updated, and maintained under both deterministic and probabilistic conditions, highlighting differences in cognitive strategies.

Numerous studies have demonstrated that mouse strains differ in neural information processing (Andolina et al., 2015b), affecting cognitive performance (Peng et al., 2023), emotional responses (Andolina et al., 2015a; Seemiller et al., 2021; Wells et al., 2026), and exploratory behavior (Andolina et al., 2015a; Seemiller et al., 2021; Sheppard et al., 2022; Wells et al., 2026). C57BL/6 mice, the most widely used laboratory strain, have been reported to show distinct sensory characteristics, including early onset progressive hearing loss (Mikaelian, 1979; Walton et al., 1995; Park et al., 2010) and congenital ocular dysfunction (Moore et al., 2018). Compared with sensorily intact CBA-derived strains, C57BL/6 mice have also been reported to exhibit alterations in synaptic plasticity (Beckmann et al., 2020; Hagena et al., 2022b) and reduced performance in hippocampus-dependent memory tasks (Kim et al., 2008; Beckmann et al., 2020). The underlying cognitive reasons for the differences in learning performance between C57BL/6 and CBA mice remain unclear. Therefore, strain-dependent variation has important implications for interpreting behavioral phenotypes, as genotype-driven differences in perception, motivation, and neural dynamics can shape how animals acquire, consolidate, and apply task rules.

The spatial T-maze is a useful paradigm for dissecting different components of associative spatial learning in rodents (Wijnen et al., 2024). When in a T-Maze, rodents naturally tend to avoid the previously visited arm, displaying spontaneous alternation behavior (Dennis, 1939; Dember and Fowler, 1958). In the associative version of the T-Maze task, however, they must overcome such a tendency to revise their choice strategy to consistently enter a single arm in order to maximize reward obtainment (André and Manahan-Vaughan, 2016; Méndez-Couz et al., 2019; Haubrich et al., 2025b). As in previous studies (Haubrich et al., 2025a), the task used here consisted of a deterministic phase followed by a probabilistic phase. In the former, reward is always present in a fixed arm (100% reward probability), allowing the animals to acquire a stable spatial rule. In the latter, the rewarded arm stays the same, but the probability of reward delivery gradually drops, while its magnitude increases to the same extent (generating correct but unrewarded outcomes, but allowing the animals to obtain the same reward size). This enables the assessment of how well animals form and maintain an established reference memory and provides insight into its perseverance and decision-making under uncertainty.

Previous work in rats has shown that distinct neural oscillatory signatures, including coordinated theta-gamma activity, track the emergence of learning strategies during T-maze learning (Rayan et al., 2022), highlighting the dynamic involvement of the hippocampus in spatial decision-making. In mice and rats, functional magnetic resonance imaging (Haubrich et al., 2025a) and in situ hybridization studies (Méndez-Couz et al., 2019; Haubrich et al., 2025b) have similarly emphasized the central role of the hippocampus and associated cortical networks in supporting learning and updating within this paradigm. Despite evidence that strain (Beckmann et al., 2020; Caliskan et al., 2022; Hagena et al., 2022a, 2022b) and age (Lester et al., 2017; Beckmann et al., 2020; Radulescu et al., 2021) influence hippocampal-cortical function, their impact on the dynamics of learning and behavioral strategies for goal-directed behavior remains unclear. We therefore investigated how strain and age influence appetitive associative spatial learning in this task, examining behavior at multiple levels, from overall task performance to trial-by-trial decision strategies.

Across all age and strain groups, correct-choice performance improved over training, consistent with effective spatial learning. However, CBA/CaOlaHsd mice reached higher levels correct performance earlier than C57BL/6 mice, indicating faster spatial learning. This advantage persisted beyond early task acquisition and remained evident when reward probability was reduced, suggesting that genetic background shapes not only how mice learn but also the stability of learned behavior when feedback becomes unreliable.

Latencies to reach goal zones provided a complementary readout of learning dynamics. Latencies decreased across training, consistent with increasing efficiency of choice execution. Older animals were slower early in training, in line with age-related reductions in response speed and/or decision speed (MacRae et al., 1988; Burwell and Gallagher, 1993; Marschner et al., 2005). CBA/CaOlaHsd mice were also faster in reaching goal zones than C57BL/6 mice during initial trials. It is not clear, however, to what extent differences in latency reflect learning and decision-making, general motivational, or motor differences.

It has long been recognized that rodents make decisions based on their past choices (Tolman, 1925). In the T-Maze paradigm, this can be studied by scrutinizing their stay/shift responses following win or loss outcomes (Salvetti et al., 2014). Accordingly, we performed a trial-by-trial analysis aimed to bridge correct-choice performance with decision policies (Smith et al., 2023; Maggi et al., 2024). In this task, the rewarded arm remained constant throughout the experiment. The probabilistic phase, therefore, created a stable but uncertain contingency environment in which occasional omissions were expected and do not indicate a reversal of the rule. Under these conditions, an adaptive policy is not to infer contingency change after an omission, but to tolerate unrewarded correct choices and continue exploiting the learned spatial rule. Already on the first training day, CBA/CaOlaHsd mice showed a higher proportion of win-stay than win-shift responses, whereas C57BL/6 mice did not show a comparable win-contingent differentiation at this early stage. However, by the end of the deterministic phase (Day 3), all groups exhibited clear asymmetries between win-stay and win-shift as well as lose-stay and lose-shift, consistent with effective learning of the task rules. In agreement with the analysis of correct choices, it shows that although all groups successfully learn the task rules, CBA/CaOlaHsd mice implemented adaptive decision policies earlier and more efficiently.

Strain and age differences became especially pronounced when reward outcomes became less reliable. During the low-probability phase, group differences emerged specifically in win-stay and in omission-driven switching after correcting but unrewarded trials. This indicates that strains differed in how omissions were incorporated into subsequent choices. In this low-probability phase, omission-driven switching predicts abandoning the correct arm after unrewarded correct choices, therefore reducing correct-choice performance. In this regard, CBA/CaOlaHsd mice appeared better adjusted to the stable but low-probability regime, maintaining exploitation despite occasional omissions. On the other hand, C57BL/6 mice showed stronger reactive updating to omissions that became disadvantageous when reward probability decreased.

In the probabilistic phase, because reward probability decreased while reward magnitude increased, there is an increase in the outcome variance, but also an increase in the payoff of sustained exploitation. The observation that higher reward magnitudes reduced between-strain differences is consistent with the idea that increasing reward value suppresses exploration by increasing the relative advantage of exploiting the known correct option. This suggests that C57BL/6 mice may require a higher reward magnitude to maintain exploitation under uncertainty, whereas CBA/CaOlaHsd mice exhibited stable exploitation even when the reward magnitude was lower.

There is a substantial body of work demonstrating strain-dependent differences in mouse hippocampal synaptic plasticity and spatial learning which are important to consider when interpreting the behavioral phenotypes described here. A well-characterized aspect of C57BL/6 mice is their early-onset, progressive sensorineural hearing loss, often beginning at high frequencies in young adulthood and worsening across months, whereas CBA/Ca-derived lines such as CBA/CaOlaHsd typically show later and milder hearing decline (Johnson et al., 1997; Spongr et al., 1997; Sha et al., 2008). Progressive hearing loss in C57BL/6 has been linked to widespread cortical adaptations and hippocampal changes, including altered receptor expression, attenuated synaptic plasticity, and impairments in spatial-memory tasks (Yu et al., 2011; Beckmann et al., 2020; Hagena et al., 2022b). Although causality between sensory decline, plasticity alterations, and learning policy cannot be established from the present behavioral data, this literature provides a strong mechanistic context for why C57BL/6 and CBA/CaOlaHsd may differ in how rapidly and efficiently spatial learning policies are encoded, stabilized, and expressed.

The older cohort (7 to 8 months) represents middle age rather than advanced aging in mice (Flurkey et al., 2007; Yanai and Endo, 2021). Accordingly, age effects on correct-choice performance were modest. Nevertheless, age influenced response latencies and omission-driven updating during the probabilistic phase. Older animals were slower early in training and showed reduced switching following losses, suggesting an age-related shift toward behavioral stability and reduced negative-outcome updating. These changes might reflect early alterations in the balance between policy flexible updating and stabilization, which may precede later declines in overall task performance (Brito et al., 2023; Attalla et al., 2024). Importantly, the consequences of age-related stabilization depended on strain-dependent strategies. Given that in C57BL/6 mice omission-driven switching was elevated during the probabilistic phase, the reduced omission-driven switching in older animals was beneficial. However, in tasks where contingencies reverse or rapid adaptation is required (Barense et al., 2002; Schoenbaum et al., 2002; Attalla et al., 2024), the same age-related reduction in switching could have been detrimental. Moreover, since here reward probability and magnitude were manipulated together, additional experiments would be needed to disentangle whether omission tolerance and strategy stabilization are driven primarily by each of these factors, and whether these factors interact with strain and age.

The reinforcement-learning analysis provided a complementary, mechanistic explanation of these strategy differences by estimating latent valence-dependent learning rates from the trial-by-trial choices. Across training, the model captured the gradual increase in expected value for the rewarded arm and the decrease for the unrewarded arm, resulting in a growing value separation that was larger in CBA/CaOlaHsd than in C57BL/6 mice, including during the probabilistic phase. Underlying this difference, CBA/CaOlaHsd mice exhibited a higher learning rate for rewarded outcomes, suggesting a faster reinforcement-driven strengthening of the correct choice policy. Moreover, an age x strain interaction in learning from unrewarded outcomes was also detected, driven by relatively higher values in younger CBA/CaOlaHsd. These learning rates were linked to observed behaviors: a higher learning rate for both rewarded and unrewarded outcomes correlated with adaptive win-stay lose-shift behavior in the deterministic phase. Importantly, only a higher learning rate for rewarded outcomes was associated with higher correct-stay and reduced omission-driven switching in the probabilistic phase. This indicates that although both learning rates help find a policy that maximizes reward when outcomes are deterministic, learning from rewarded trials is more important when the outcomes become noisier. Therefore, strain differences in performance under uncertainty seem to reflect differences in how rapidly and strongly reward information is incorporated and translated into actions.

In summary, both CBA/CaOlaHsd and C57BL/6 learned a spatial appetitive memory task, but differed as to how they reached and maintained high correct-choice performance. CBA/CaOlaHsd mice reached high correct-choice performance earlier and expressed a policy that maintained exploitation when reward delivery became unreliable. In contrast, C57BL/6 mice were more prone to omission-driven switching, a strategy that is not optimal when the correct option remains constant, but outcomes are probabilistic. These differences were consistent with stronger learning rates from rewarded outcomes in CBA/CaOlaHsd. Age mainly affected latencies and reduced omission-driven switching, indicating an age-related shift toward greater behavioral stability. Together, these findings highlight that genetic background and age shape not only learning success but also underlying decision policies and learning dynamics. Future studies shall further scrutinize the strain- and age-specific physiology underlying these effects. Beyond their mechanistic implications, these findings have also practical relevance for murine strain selection in neuroscience research. For instance, our data suggests that CBA/CaOlaHsd mice may be particularly suitable for studies investigating reinforcement-driven spatial learning supported by stable reference memory, whereas C57BL/6 mice may be especially informative for examining sensitivity to immediate feedback and outcome fluctuations. Given these strain-dependent differences in reward-driven learning, sensitivity to uncertainty, and stability of learned spatial preferences, careful consideration is warranted when interpreting and comparing behavioral outcomes across mouse strains.

## Conflict of Interest

*The authors declare that the research was conducted in the absence of any commercial or financial relationships that could be construed as a potential conflict of interest*.

## Author Contributions

The study was designed by D.M.-V. Experiments and behavioral task performance analysis were conducted by J.L. and J.H. Analysis of trial-by-trial learning, as well as the development and implementation of the reinforcement learning model were conducted by J.H. Data were interpreted and the paper was written by all authors.

## Funding

This work was supported by a German Research Foundation (Deutsche Forschungsgemeinschaft) grant to D.M.-V (SFB 1280/A04, project number: 316803389). J.L. is a recipient of a stipend from the Chinese Scholarship Council.

## Acknowledgments

We are grateful to Dr. Thu-Huong Hoang for technical advice, and Juliane Böge and Jens Colitti-Klausnitzer for technical assistance with animal treatments. We thank Nadine Kollosch for animal care.

## Notes

### Competing Interest Statement

The authors have declared no competing interest.

